# Alpha-Synuclein Co-Pathology and Amyloid–Tau Coupling in Alzheimer’s Disease: Cross-Sectional and Longitudinal Analyses Across Braak Stage Regions

**DOI:** 10.64898/2026.03.31.713304

**Authors:** Ahmed Negida, Jamir Pitton Rissardo, Alzheimer’s Disease Neuroimaging Initiative

**Affiliations:** Department of Neurology, Virginia Commonwealth University, Richmond, VA, USA; Parkinson’s and Movement Disorders Center, Virginia Commonwealth University, Richmond, VA, USA; Neurology Department, Cooper University Hospital, Camden, NJ 08103, USA

**Keywords:** Alzheimer’s disease, alpha-synuclein, seed amplification assay, tau PET, Lewy body, longitudinal

## Abstract

**Background:** Misfolded alpha-synuclein (αSyn), a common co-pathology in Alzheimer’s disease, acts synergistically with amyloid and tau in animal models, but its role in the human amyloid–tau–cognition cascade is unclear. We examined whether αSyn modifies the amyloid–tau and tau–cognition associations, cross-sectionally and over time.

**Methods:** We analyzed 615 participants from the Alzheimer’s Disease Neuroimaging Initiative with a cerebrospinal fluid αSyn seed amplification assay (SAA), flortaucipir tau positron emission tomography (PET) in Braak stage regions, and amyloid PET and domain-specific cognitive composites within 1 year of tau PET. Linear models with heteroscedasticity-consistent standard errors tested αSyn × amyloid interactions on regional tau and αSyn × tau interactions on cognition; mixed-effects models tested the same interactions on repeated tau PET (n = 341) and cognitive testing (n = 516).

**Results:** αSyn positivity was present in 117 participants (19.0%). Cross-sectionally, the amyloid–tau association was steeper in αSyn SAA+ participants for meta-temporal tau (interaction β 0.258, 95% CI 0.023 to 0.493; p = 0.032) and Braak I–IV (p ≤ 0.048); no interaction survived false discovery rate correction (q = 0.072), and clinical-stage adjustment attenuated it (β 0.114; p = 0.333). Over a median of 3.5 years (n = 341; 68 αSyn-positive), no αSyn × amyloid × time interaction was detected (meta-temporal β −0.012, 95% CI −0.046 to 0.023; p = 0.513). No cross-sectional αSyn × tau interaction on cognition was detected; at mean tau, αSyn positivity was associated with lower executive function (q = 0.032). Longitudinally, tau-related executive decline was less steep in αSyn-positive participants (q = 0.042).

**Conclusions:** The αSyn-associated amyloid–tau interaction was modest, did not withstand multiple-testing correction, and had no detected longitudinal counterpart. αSyn positivity was associated with poorer executive function at mean tau; the unexpected longitudinal executive interaction needs replication. These findings do not establish αSyn status as a marker for tau-progression enrichment in trials.

## 1. Introduction

Amyloid-β (Aβ) deposition is common in older adults without dementia and marks the start of the Alzheimer’s disease (AD) continuum in biological definitions of the disease [1, 2]. However, individuals with similar amyloid burden can follow different clinical trajectories [3, 4], suggesting that other factors contribute to disease progression. The NIA-AA amyloid-tau-neurodegeneration (ATN) framework places Aβ upstream of tau pathology, neurodegeneration, and cognitive decline [1]. Identifying the biological modifiers that accelerate or slow specific steps in this cascade matters for prognosis, for trial design, and for the timing of anti-amyloid therapy.

Alpha-synuclein (αSyn) co-pathology is a leading candidate modifier because it is both frequent and biologically plausible. Lewy body pathology is found in approximately 40–60% of autopsy-confirmed AD cases, making it the most common co-pathology in the disease [5, 6]. Experimental work links αSyn to both Aβ and tau: in transgenic mice, Aβ, tau, and αSyn interact synergistically and accelerate neuropathology and cognitive decline beyond what any single proteinopathy produces [7]; distinct αSyn fibril strains cross-seed tau aggregation in primary neurons and in vivo [8]; and soluble αSyn oligomers alter the aggregation kinetics of Aβ and tau [9, 10]. Whether these interactions operate in living patients, and at which step of the cascade, remains unclear.

Until recently, αSyn in AD could be studied only at autopsy. Cerebrospinal fluid (CSF) seed amplification assays (SAA) now detect misfolded αSyn in vivo with high sensitivity and specificity, as shown at scale in the Parkinson’s Progression Markers Initiative (PPMI) [11]. In the Alzheimer’s Disease Neuroimaging Initiative (ADNI), the assay detected autopsy-confirmed neocortical Lewy pathology in every case but limbic and amygdala-predominant pathology less reliably [12], and SAA positivity was associated with greater amyloid burden, earlier symptom onset, and faster cognitive decline [13]. Across ADNI and BioFINDER-2, αSyn positivity tracked Aβ rather than tau [14], and in clinically unimpaired adults it had cognitive effects independent of AD pathology [15].

Franzmeier et al. addressed the amyloid-to-tau question directly in 592 ADNI participants [16]. In αSyn SAA-positive participants, amyloid PET was more steeply associated with tau PET in the temporal meta-region and in Braak stage regions, and in the 290 participants with repeated scans over about 2.5 years, amyloid-related tau accumulation was faster. The authors also reported faster amyloid-related decline on the 13-item Alzheimer’s Disease Assessment Scale–Cognitive Subscale but no αSyn × tau PET interaction on cognitive change. Two questions remain open. First, it is unknown whether the longitudinal acceleration holds as tau PET follow-up in ADNI lengthens. Second, a global cognitive scale may miss domain-specific effects: Lewy pathology without AD co-pathology produces a predominantly dysexecutive and visuospatial profile, whereas mixed AD and Lewy pathology remains memory-predominant [17].

Here we re-examined these relationships in ADNI participants with a CSF αSyn SAA result, amyloid PET, regional tau PET in Braak stage regions, structural MRI, and domain-specific cognitive assessments, matching amyloid PET and cognition to the tau PET scan and adding all tau PET and cognitive follow-up available in January 2026. We asked whether αSyn positivity (1) modifies the cross-sectional association between amyloid and tau across Braak stage regions; (2) modifies the rate of amyloid-related tau accumulation over time; and (3) modifies the cross-sectional or longitudinal association between tau and memory, executive function, language, and visuospatial function.

## 2. Methods

### Study design and participants

Data were obtained from the ADNI database (adni.loni.usc.edu). The ADNI was launched in 2003 as a public-private partnership, led by Principal Investigator Michael W. Weiner, MD. The primary goal of ADNI has been to test whether serial MRI, PET, other biological markers, and clinical and neuropsychological assessment can be combined to measure the progression of mild cognitive impairment (MCI) and early AD. For up-to-date information, see www.adni-info.org [18]. Participants included cognitively normal (CN), MCI, and dementia participants enrolled in ADNI-1, ADNI-GO, ADNI-2, and ADNI-3. The ADNI study was approved by the Institutional Review Board or Independent Ethics Committee at each participating site, and all participants provided written informed consent in accordance with the Declaration of Helsinki.

### Inclusion and exclusion criteria

We included ADNI participants with a determinate CSF αSyn SAA result, at least 1 amyloid PET scan, at least 1 flortaucipir tau PET scan, a T1-weighted MRI processed through the MUSE pipeline, PHC cognitive composite scores, and APOE genotype. Of 1,658 SAA results, 9 were indeterminate and were excluded; the remaining 1,649 results came from 1,629 unique participants, and for the 20 participants with more than 1 result (all concordant) the earliest was used. Of these 1,629 participants, 1,350 had amyloid PET, 663 also had tau PET, and 657 also had MUSE-processed MRI; cognitive composites and APOE genotype were available for all 657. After exclusion of 21 participants with a missing covariate or outcome, the analytical cohort comprised 636 participants (Figure 1). Modalities were not required to come from the same visit. For the primary cross-sectional analyses, amyloid PET and cognitive composites were taken from the examination nearest baseline tau PET, and participants with either modality more than 1 year from tau PET were excluded (n = 21), giving a primary sample of 615. MRI came from the earliest quality-controlled examination. Age was taken at baseline tau PET, and clinical diagnosis (CN, MCI, or dementia) from the ADNI diagnostic summary at the visit nearest baseline tau PET (median interval 15 days, IQR 6 to 42; ties resolved toward the earlier visit). Analyses that used the earliest examination of every modality in all 636 participants are reported as a sensitivity analysis. Intervals between modalities are reported in Supplementary Table S5.

**Figure 1.**
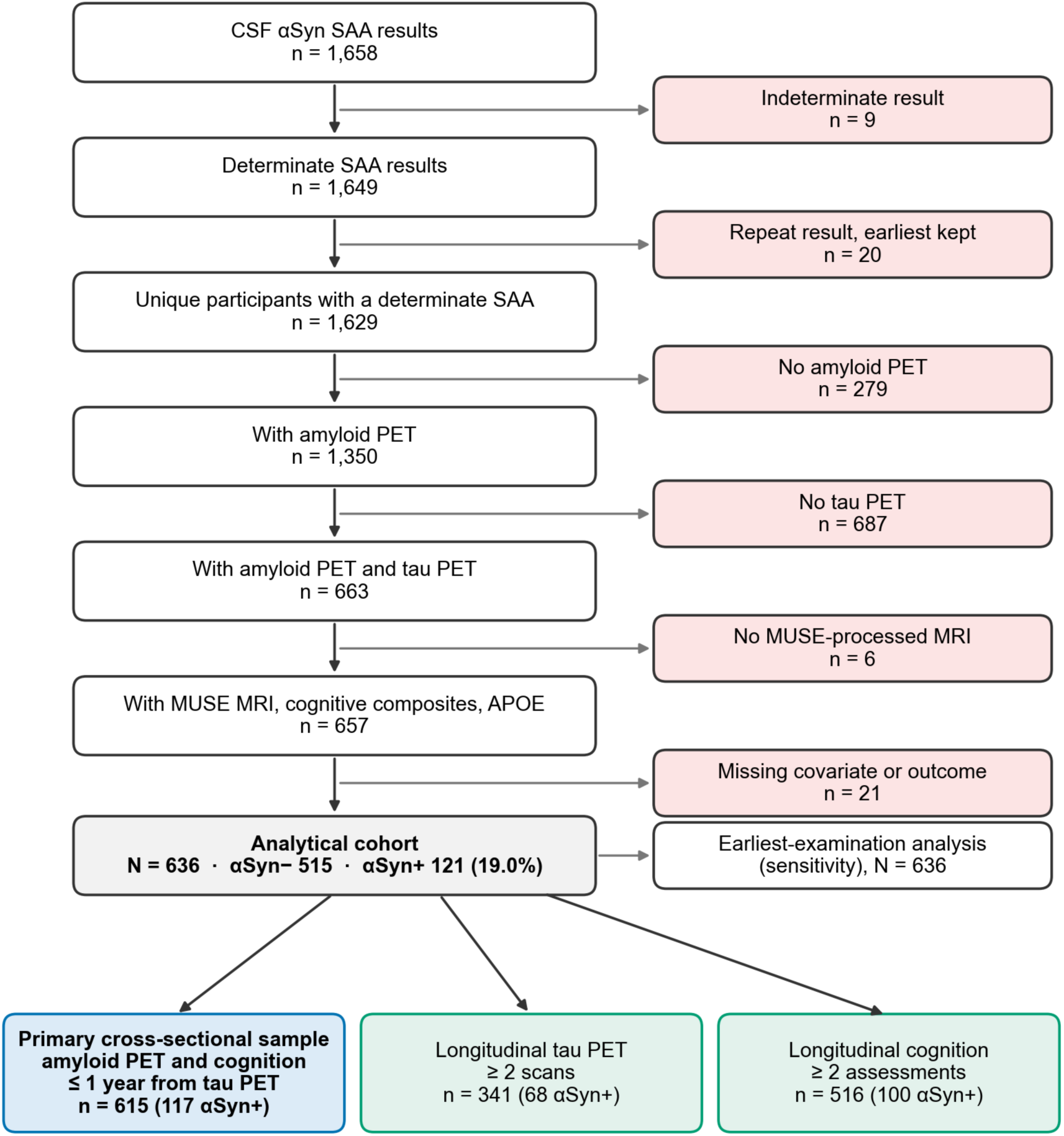
Study flow diagram. Of 1,658 cerebrospinal fluid alpha-synuclein seed amplification assay (SAA) results, 1,649 were determinate and came from 1,629 unique participants; 636 of these had amyloid PET, tau PET, MUSE-processed MRI, cognitive composites, APOE genotype, and complete covariates (analytical cohort). Of the 636, 615 had amyloid PET and cognitive testing within 1 year of baseline tau PET and formed the primary cross-sectional sample, 341 had at least 2 tau PET scans, and 516 had at least 2 cognitive assessments; the earliest examinations of all 636 were used in a sensitivity analysis. Abbreviations: αSyn, alpha-synuclein; APOE, apolipoprotein E; MRI, magnetic resonance imaging; MUSE, Multi-atlas region Segmentation utilizing Ensembles; PET, positron emission tomography.

### Alpha-synuclein seed amplification assay

αSyn aggregation was assessed using the Amprion CSF-based SAA, which detects misfolded αSyn seeds through cyclic amplification of recombinant αSyn substrate [11]. The assay reports Detected-1 (amplification profile typical of Parkinson’s disease or dementia with Lewy bodies), Detected-2 (profile typical of multiple system atrophy), Not Detected, or Indeterminate. Participants with Detected-1 or Detected-2 results were classified as αSyn-positive (primary sample: n = 115 and n = 2, respectively) and those with Not Detected results as αSyn-negative; indeterminate results were excluded [13]. SAA status was based on a single CSF sample, timed by its collection date rather than its assay date, and was treated as time-invariant. In the primary sample, CSF was collected a median of 1.73 years (IQR 0.04 to 3.72) from baseline tau PET: on or before the tau PET date in 291 participants (47.3%) and more than 1 year after it in 204 (33.2%).

### Amyloid PET

Amyloid PET was performed using florbetapir ([18F]AV-45; n = 398) or florbetaben ([18F]FBB; n = 217) for the scans used in the primary sample, with centiloid values computed through the Alzheimer’s Disease Sequencing Project Phenotype Harmonization Consortium (PHC) pipeline to enable cross-tracer comparability [19]. Amyloid positivity was defined by the PHC tracer-specific amyloid status of the same scan. The primary cross-sectional and the longitudinal analyses used the quality-controlled scan nearest baseline tau PET; the earliest-examination sensitivity analysis used each participant’s earliest scan (florbetapir, n = 412; florbetaben, n = 216; Pittsburgh Compound B, n = 8).

### Tau PET

Tau PET was performed using flortaucipir ([18F]AV-1451). Regional standardized uptake value ratios (SUVRs) were extracted from the ADNI PHC detailed tau dataset, which provides Braak stage-specific composites (Braak I–II, III, IV, V, VI) and a meta-temporal composite [20]. Tau positivity was defined as meta-temporal SUVR > 1.32, the conservative cut point reported by Jack et al., derived from the accuracy of discriminating amyloid-positive cognitively impaired from age- and sex-matched amyloid-negative clinically normal individuals [21]. Because this cut point was derived with a different processing pipeline, tau status was used only for descriptive AT grouping and not in the primary interaction models. Scans failing quality control were excluded. The earliest scan served as baseline, and all later scans were used for the longitudinal analyses.

### Structural MRI and neurodegeneration markers

T1-weighted structural MRI scans were processed using the MUSE (Multi-atlas region Segmentation utilizing Ensembles) pipeline [22]. Hippocampal volume was calculated as the sum of left and right hippocampal MUSE ROIs (ROIs 47 and 48). The SPARE-AD (Spatial Pattern of Abnormality for Recognition of Early AD) score, an MRI-based machine-learning index of AD-typical neurodegeneration, was used to define neurodegeneration status (N+) using a cohort-median split [23]. Higher SPARE-AD values indicate more AD-like brain patterns.

### Cognitive assessment

Cognitive performance was assessed using the ADNI PHC cognitive composite scores, which provide domain-specific measures for memory (PHC_MEM), executive function (PHC_EXF), language (PHC_LAN), and visuospatial processing (PHC_VSP) [24]. These composites are derived from multiple neuropsychological tests using item response theory and are optimized for sensitivity to change across the AD spectrum. The primary analyses used the assessment nearest baseline tau PET.

### ATN classification

Participants were classified according to the NIA-AA ATN framework [1]. Amyloid (A) status was determined by the PHC tracer-specific status of the amyloid PET scan nearest tau PET. Tau (T) status was defined by meta-temporal SUVR > 1.32. Neurodegeneration (N) status was defined by a SPARE-AD score above the median of the 657 participants with MUSE-processed MRI (the cohort before the exclusion of 21 participants with a missing covariate or outcome). This produced 8 ATN profiles (A±, T±, N±) and 4 simplified AT groups (A−T−, A−T+, A+T−, A+T+).

### Statistical analysis

Baseline characteristics were compared between αSyn-positive and αSyn-negative groups using Mann–Whitney U tests (continuous variables) and chi-square tests (categorical variables). αSyn positivity prevalence across AT groups was compared using the chi-square test of independence.

Primary analyses used ordinary least squares (OLS) linear regression with interaction terms in the primary sample (n = 615) to test 2 hypotheses. First, whether αSyn modifies the amyloid–tau association: tau_z ∼ centiloids_z × αSyn_pos + age_z + sex + education_z + APOE4. Six models were fit, one for each tau region (meta-temporal, Braak I–II, III, IV, V, VI). Second, whether αSyn modifies the tau–cognition association: cognition_z ∼ meta-temporal_tau_z × αSyn_pos + centiloids_z + age_z + sex + education_z + APOE4. Four models were fit, one for each cognitive domain. Continuous predictors and outcomes were standardized with the mean and standard deviation of the primary sample; the amyloid-positive, diagnosis-stratified, CSF-restricted, and Detected-2-excluded sensitivity analyses kept these reference values, so their coefficients are on the primary scale, whereas the earliest-examination analysis and the models adjusted for diagnosis and severity were standardized within their own samples. Because residual variance increased with amyloid and tau burden (Breusch–Pagan p < 0.001 in all 6 amyloid-to-tau models), all cross-sectional inferences used heteroscedasticity-consistent (HC3) standard errors, with two-sided t tests and confidence intervals on the residual degrees of freedom [25]. Significance was set at α = 0.05 (two-sided). Given the hypothesis-driven nature of the analysis, no correction for multiple comparisons was applied to the primary interaction tests [26]. As a sensitivity analysis, false discovery rate (FDR) correction using the Benjamini-Hochberg procedure was applied across the 6 amyloid-to-tau interaction tests. As a supplementary analysis added at a coauthor’s request, the tau–cognition interaction models were repeated using Braak stage-specific tau PET composites (Braak I–II, III, IV, V, and VI) in place of meta-temporal tau to determine whether regional tau burden influenced the observed findings.

Sensitivity analyses (1) to (4) were planned with the primary analysis: (1) restriction to amyloid-positive participants; (2) stratification by clinical diagnosis (CN and MCI; the dementia group, n = 66, was too small to analyze separately); (3) logistic regression models for AT group membership; and (4) αSyn main-effect models without the interaction term. Analysis (5), added at a coauthor’s request, was additional adjustment of the primary interaction models for clinical diagnosis (CN, MCI, dementia), Clinical Dementia Rating Sum of Boxes (CDR-SB), and Mini-Mental State Examination (MMSE) score, with diagnosis, CDR-SB, and MMSE taken from the assessment nearest each participant’s tau PET scan. Three further sensitivity analyses were added during revision: (6) the earliest available examination of every modality in all 636 participants; (7) restriction to participants whose CSF sample was within 2 years of tau PET (n = 356); and (8) exclusion of the 2 participants with a Detected-2 result. The analysis plan was also revised after the first submission: the primary sample was changed from the earliest examination of every modality to examinations matched to tau PET, HC3 standard errors replaced conventional ones, the tau cut point was changed from 1.30 to the published value of 1.32, and age and diagnosis, previously taken from the first amyloid PET visit, were taken at tau PET; results under the earlier choices are reported in the text and in Supplementary Tables S6 and S10. All analyses were conducted in Python 3.13 (statsmodels 0.14.5, scipy 1.16, pandas 2.3, numpy 2.3), with the random seed set to 2025.

### Longitudinal analyses

All flortaucipir scans acquired on or after baseline tau PET were extracted from the PHC tau dataset, and 341 participants had at least 2 scans. Regional SUVRs were divided by the baseline cohort standard deviation so that slopes are expressed in baseline SD per year. Linear mixed-effects models with random intercepts and random slopes for time (years from baseline tau PET) were fitted by restricted maximum likelihood: tau ∼ time × centiloids_z × αSyn_pos + time × (age_z + sex + education_z + APOE4), with all lower-order terms and with centiloids from the amyloid PET scan nearest baseline tau PET; continuous covariates were standardized in the 636-participant analytical cohort, with age at baseline tau PET. Each model was fitted with 4 optimization algorithms, and the solution with the highest restricted likelihood was retained. The time × centiloids × αSyn term tested whether αSyn modified amyloid-related tau accumulation, and the time × αSyn term tested whether αSyn-positive participants accumulated tau faster at average amyloid burden; model-based group differences in annual meta-temporal accumulation were also estimated at 0, 50, and 100 centiloids. The models were refit in amyloid-positive participants (PHC tracer-specific status of the scan nearest baseline tau PET, as in the cross-sectional analyses, and, as an alternative threshold, ≥20 centiloids), with additional adjustment for diagnosis at baseline tau PET, and, post hoc, in participants whose CSF sample was collected no later than 1 year after baseline tau PET or within 2 years of it.

For cognition, all PHC composite assessments from 6 months before baseline tau PET onward were used in participants with at least 2 assessments (n = 516). Models included time × baseline meta-temporal tau_z × αSyn_pos and time interactions with centiloids, age, sex, education, and APOE4, with random intercepts and slopes. The Benjamini-Hochberg procedure was applied within each family of 3-way interaction tests (6 tau regions; 4 cognitive domains). To assess attrition bias, we compared participants with and without tau PET follow-up and refit the tau model as a 2-stage analysis of participant-specific slopes weighted by the inverse probability of follow-up, estimated from age, sex, education, APOE4, centiloids, αSyn status, and diagnosis. The number of participants observed at each year of follow-up is reported in Supplementary Table S9.

Two post hoc analyses addressed published ADNI findings. First, to test whether the analytic approach explained the difference from Franzmeier et al. [16], we refit the tau models in the same participants and scans as a 2-stage analysis like theirs: participant-specific annual change was predicted from a mixed model with random intercepts and slopes and no covariates, then regressed on centiloids × αSyn with follow-up time and either our covariates or theirs (age, sex, and study site), all standardized. Unlike the 2-stage approach, a 1-stage mixed model estimates change and its predictors jointly and uses every scan [27]. After seeing the 2-stage result, we added 2 diagnostics: the second stage was also adjusted for baseline tau, and the predicted rates were replaced by each participant’s least-squares slope. Second, because faster tau accumulation has been reported in SAA-positive women [28], we fitted an exploratory time × αSyn × sex model with all lower-order terms.

## 3. Results

### Cohort characteristics

Of 1,629 ADNI participants with a determinate αSyn SAA result, 636 met inclusion criteria after requiring amyloid PET, tau PET, structural MRI, cognitive composites, and APOE genotyping, and 615 had amyloid PET and cognitive testing within 1 year of baseline tau PET and formed the primary cross-sectional sample (Figure 1). At baseline tau PET, the primary sample comprised 349 CN (56.7%), 200 MCI (32.5%), and 66 dementia (10.7%) participants, with a mean age of 73.6 years (SD 7.8), 52.2% female, and mean education of 16.5 years (SD 2.4). Amyloid PET and cognitive testing were a median of 0.03 and 0.04 years from tau PET; MRI was more than 1 year from tau PET in 252 participants (41.0%) (Supplementary Table S5).

αSyn SAA positivity was detected in 117 participants (19.0%). Compared with αSyn-negative participants, αSyn-positive participants were older (mean 75.7 vs. 73.1 years; p < 0.001) and had higher amyloid burden (39.7 vs. 27.5 centiloids; p = 0.008), higher meta-temporal tau SUVR (1.44 vs. 1.31; p < 0.001), higher SPARE-AD scores (−0.08 vs. −0.36; p < 0.001), smaller hippocampal volume (7,066 vs. 7,361 mm³; p = 0.006), and lower memory (0.22 vs. 0.59; p < 0.001), executive function (0.32 vs. 0.68; p < 0.001), and language (0.39 vs. 0.67; p < 0.001) composite scores. APOE ε4 carriage was more frequent in the αSyn-positive group (47.0% vs. 36.3%; p = 0.042), as was dementia (24.8% vs. 7.4%; diagnosis distribution p < 0.001) (Table 1). Because the groups differed in clinical stage, we also fitted models adjusted for diagnosis and severity (see Sensitivity analyses).

**Table 1.**
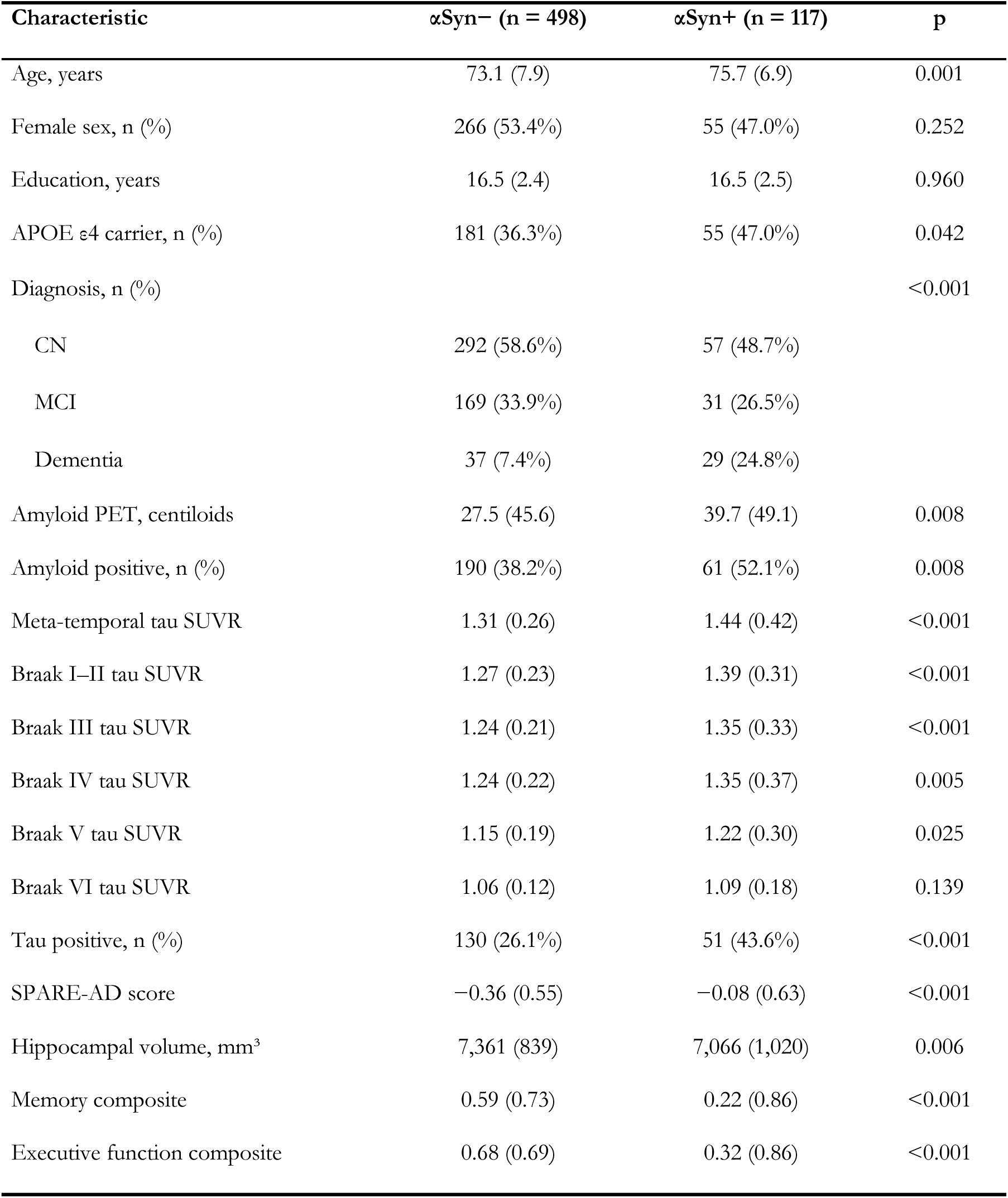
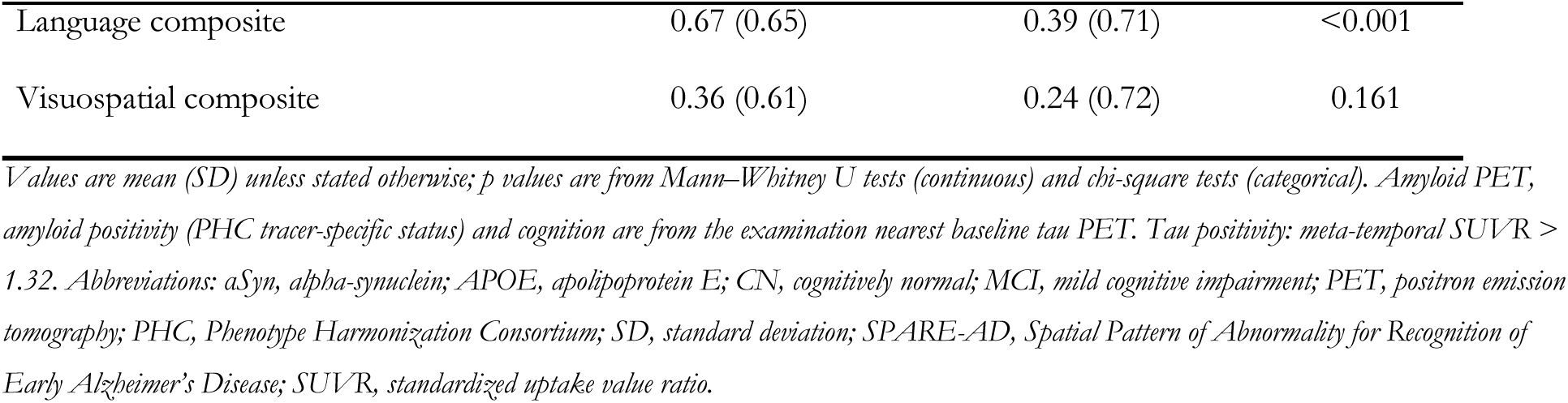
Baseline characteristics of the primary sample by αSyn SAA status (n = 615). Continuous variables are mean (SD) and were compared with Mann–Whitney U tests; categorical variables are n (%) and were compared with chi-square tests. Amyloid PET and cognition are from the examination nearest baseline tau PET; age and diagnosis are at baseline tau PET.

| Characteristic | αSyn– (n = 498) | αSyn+ (n = 117) | p |
| --- | --- | --- | --- |
| Age, years | 73.1 (7.9) | 75.7 (6.9) | 0.001 |
| Female sex, n (%) | 266 (53.4%) | 55 (47.0%) | 0.252 |
| Education, years | 16.5 (2.4) | 16.5 (2.5) | 0.960 |
| APOE ε4 carrier, n (%) | 181 (36.3%) | 55 (47.0%) | 0.042 |
| Diagnosis, n (%) |  |  | <0.001 |
| CN | 292 (58.6%) | 57 (48.7%) |  |
| MCI | 169 (33.9%) | 31 (26.5%) |  |
| Dementia | 37 (7.4%) | 29 (24.8%) |  |
| Amyloid PET, centiloids | 27.5 (45.6) | 39.7 (49.1) | 0.008 |
| Amyloid positive, n (%) | 190 (38.2%) | 61 (52.1%) | 0.008 |
| Meta-temporal tau SUVR | 1.31 (0.26) | 1.44 (0.42) | <0.001 |
| Braak I–II tau SUVR | 1.27 (0.23) | 1.39 (0.31) | <0.001 |
| Braak III tau SUVR | 1.24 (0.21) | 1.35 (0.33) | <0.001 |
| Braak IV tau SUVR | 1.24 (0.22) | 1.35 (0.37) | 0.005 |
| Braak V tau SUVR | 1.15 (0.19) | 1.22 (0.30) | 0.025 |
| Braak VI tau SUVR | 1.06 (0.12) | 1.09 (0.18) | 0.139 |
| Tau positive, n (%) | 130 (26.1%) | 51 (43.6%) | <0.001 |
| SPARE-AD score | −0.36 (0.55) | −0.08 (0.63) | <0.001 |
| Hippocampal volume, mm<sup>3</sup> | 7,361 (839) | 7,066 (1,020) | 0.006 |
| Memory composite | 0.59 (0.73) | 0.22 (0.86) | <0.001 |
| Executive function composite | 0.68 (0.69) | 0.32 (0.86) | <0.001 |

| Characteristic | αSyn$-$ (n = 498) | αSyn$+$ (n = 117) | p |
| --- | --- | --- | --- |
| Language composite | 0.67 (0.65) | 0.39 (0.71) | <0.001 |
| Visuospatial composite | 0.36 (0.61) | 0.24 (0.72) | 0.161 |

Values are mean (SD) unless stated otherwise; p values are from Mann–Whitney U tests (continuous) and chi-square tests (categorical). Amyloid PET, amyloid positivity (PHC tracer-specific status) and cognition are from the examination nearest baseline tau PET. Tau positivity: meta-temporal SUVR > 1.32. Abbreviations: αSyn, alpha-synuclein; APOE, apolipoprotein E; CN, cognitively normal; MCI, mild cognitive impairment; PET, positron emission tomography; PHC, Phenotype Harmonization Consortium; SD, standard deviation; SPARE-AD, Spatial Pattern of Abnormality for Recognition of Early Alzheimer’s Disease; SUVR, standardized uptake value ratio.

### αSyn positivity across AT biomarker profiles

The prevalence of αSyn SAA positivity differed across AT biomarker groups (χ² = 15.07, df = 3, p = 0.002). It was highest in the A−T+ (29.0%; 9 of 31) and A+T+ (28.0%; 42 of 150) groups, followed by A+T− (18.8%; 19 of 101) and A−T− (14.1%; 47 of 333). Within the full ATN framework, αSyn positivity ranged from 11.4% in A−T−N− (23 of 202) to 43.8% in A−T+N+ (7 of 16); among groups with at least 30 participants, it was highest in A+T+N+ (32.4%; 36 of 111) (Table 2, Figure 2).

**Table 2.**
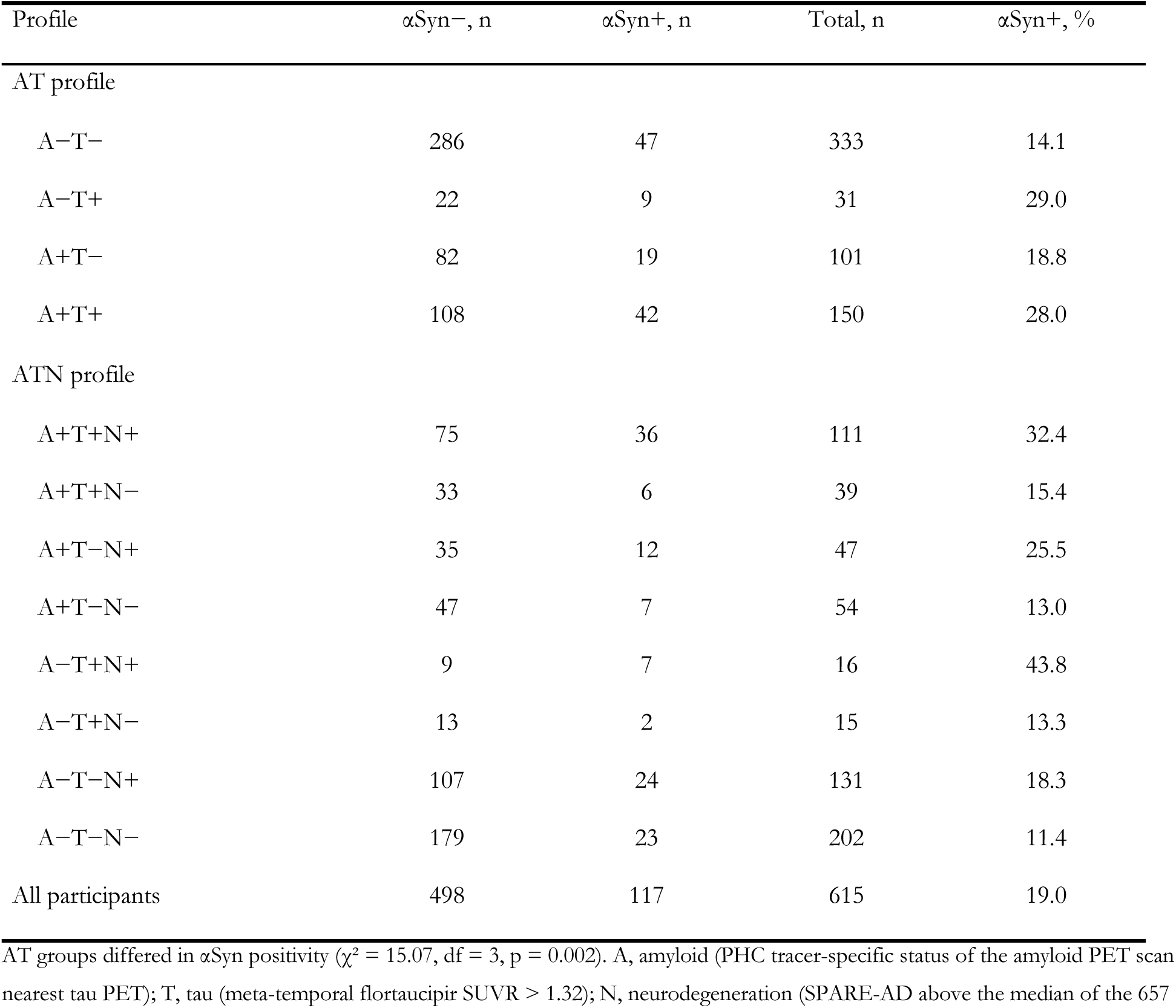
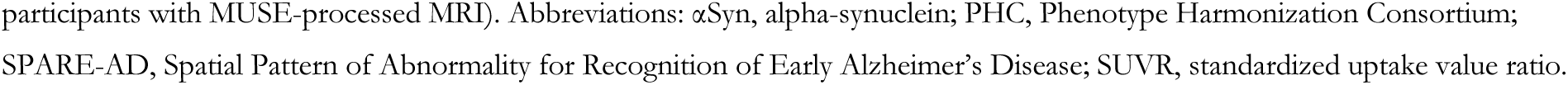
Prevalence of αSyn SAA positivity across AT and ATN biomarker profiles (n = 615). A, amyloid (PHC tracer-specific status of the amyloid PET scan nearest tau PET); T, tau (meta-temporal flortaucipir SUVR > 1.32); N, neurodegeneration (SPARE-AD above the median of the 657 participants with MUSE-processed MRI).

**Figure 2.**
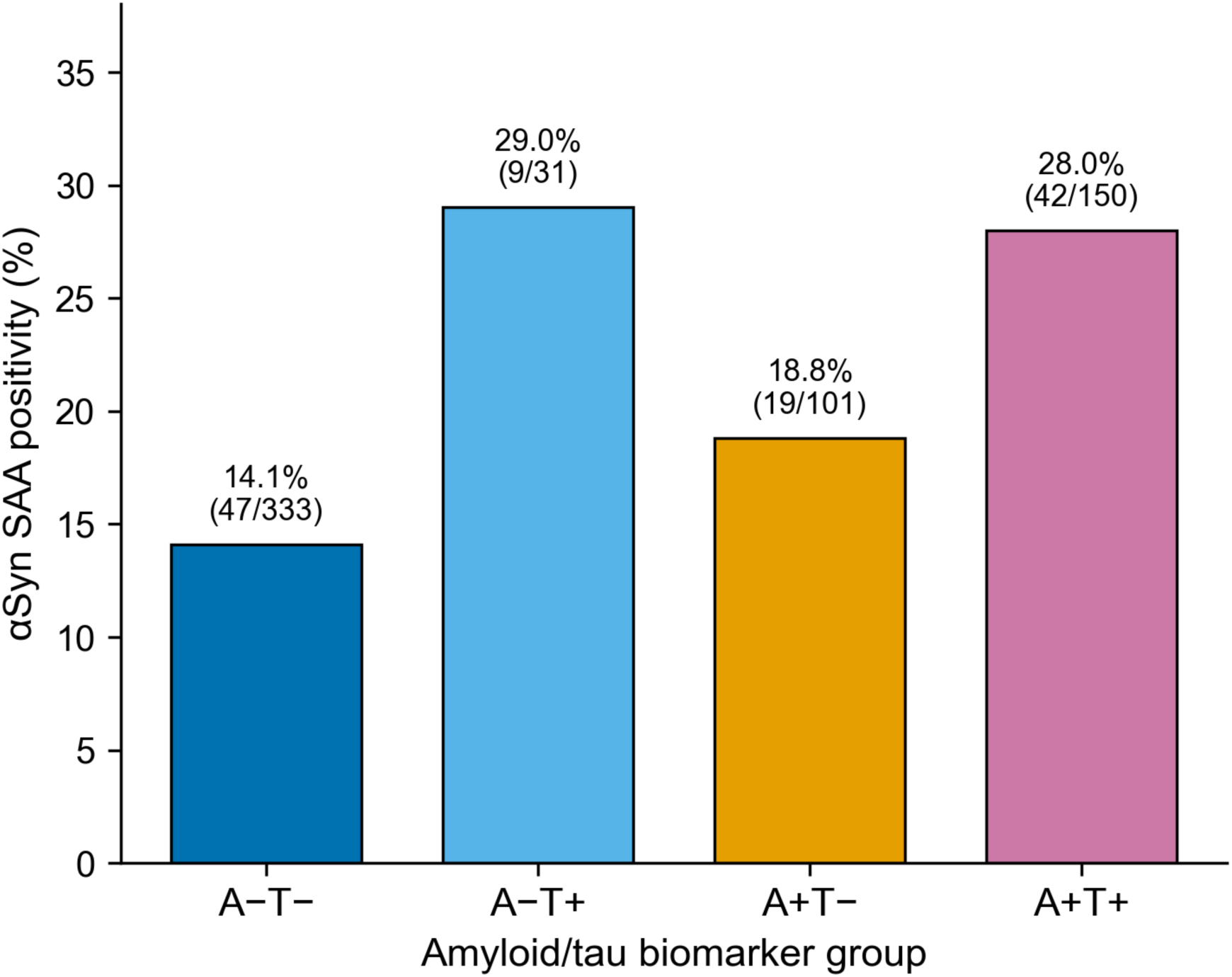
Prevalence of αSyn SAA positivity across AT biomarker groups (n = 615). Bars show the percentage of αSyn-positive participants in each group, with percentages and counts above the bars; groups were compared with a chi-square test (χ² = 15.07, df = 3, p = 0.002). A, amyloid (scan nearest tau PET); T, tau (meta-temporal SUVR > 1.32). Abbreviations: αSyn, alpha-synuclein; SAA, seed amplification assay; SUVR, standardized uptake value ratio.

### Cross-sectional amyloid–tau coupling by αSyn status

In models adjusted for age, sex, education, and APOE ε4 status, higher amyloid burden (centiloids) was associated with higher tau PET uptake in every region (all p < 0.001). The association was steeper in αSyn-positive participants for meta-temporal tau (αSyn × centiloids β = 0.258, 95% CI 0.023 to 0.493, p = 0.032), Braak I to II (β = 0.196, 95% CI 0.003 to 0.390, p = 0.047), Braak III (β = 0.262, 95% CI 0.007 to 0.516, p = 0.044), and Braak IV (β = 0.253, 95% CI 0.002 to 0.504, p = 0.048), but not for Braak V (β = 0.217, 95% CI −0.102 to 0.537, p = 0.182) or Braak VI (β = 0.189, 95% CI −0.183 to 0.561, p = 0.319) (Table 3, Figure 3). The interaction estimates were 38% to 57% of the amyloid main effect. None of the 6 interactions survived FDR correction across the regional models (q = 0.072 for meta-temporal and Braak I–IV tau; Supplementary Table S1). With conventional OLS standard errors, all 6 interactions would have been significant (p = 0.002 to 0.034); the difference reflects greater residual variance at high amyloid and tau burden.

**Table 3.** Cross-sectional αSyn × amyloid interaction on regional tau PET (n = 615). Each row is a linear regression of z-standardized regional tau SUVR on z-standardized centiloids, αSyn status, and their interaction, adjusted for age, sex, education, and APOE ε4 carrier status. β values are standardized coefficients; p values and 95% CIs use heteroscedasticity-consistent (HC3) standard errors, and CIs are shown for the interaction terms.

| Tau region | $\beta$ amyloid | $\beta$ αSyn | $\beta$ interaction (95% CI) | p interaction | Adj. R<sup>2</sup> |
| --- | --- | --- | --- | --- | --- |
| Meta-temporal | 0.470 (p < 0.001) | 0.275 (p = 0.011) | 0.258 (0.023, 0.493) | 0.032 | 0.318 |
| Braak I–II | 0.512 (p < 0.001) | 0.267 (p = 0.004) | 0.196 (0.003, 0.390) | 0.047 | 0.391 |
| Braak III | 0.463 (p < 0.001) | 0.296 (p = 0.005) | 0.262 (0.007, 0.516) | 0.044 | 0.313 |
| Braak IV | 0.452 (p < 0.001) | 0.274 (p = 0.016) | 0.253 (0.002, 0.504) | 0.048 | 0.296 |
| Braak V | 0.444 (p < 0.001) | 0.233 (p = 0.019) | 0.217 (−0.102, 0.537) | 0.182 | 0.269 |
| Braak VI | 0.406 (p < 0.001) | 0.197 (p = 0.052) | 0.189 (−0.183, 0.561) | 0.319 | 0.201 |

$\beta$ values are standardized coefficients from linear regression of z-scored regional tau SUVR on z-scored centiloids, αSyn status and their interaction, adjusted for age, sex, education and APOE ε4 carrier status (n = 615); p values and 95% CIs use heteroscedasticity-consistent (HC3) standard errors. $\beta$ amyloid is the centiloid slope in αSyn-negative participants; $\beta$ αSyn is the αSyn difference at mean centiloids. Abbreviations: αSyn, alpha-synuclein; Adj., adjusted; APOE, apolipoprotein E; CI, confidence interval; SUVR, standardized uptake value ratio.

**Figure 3.**
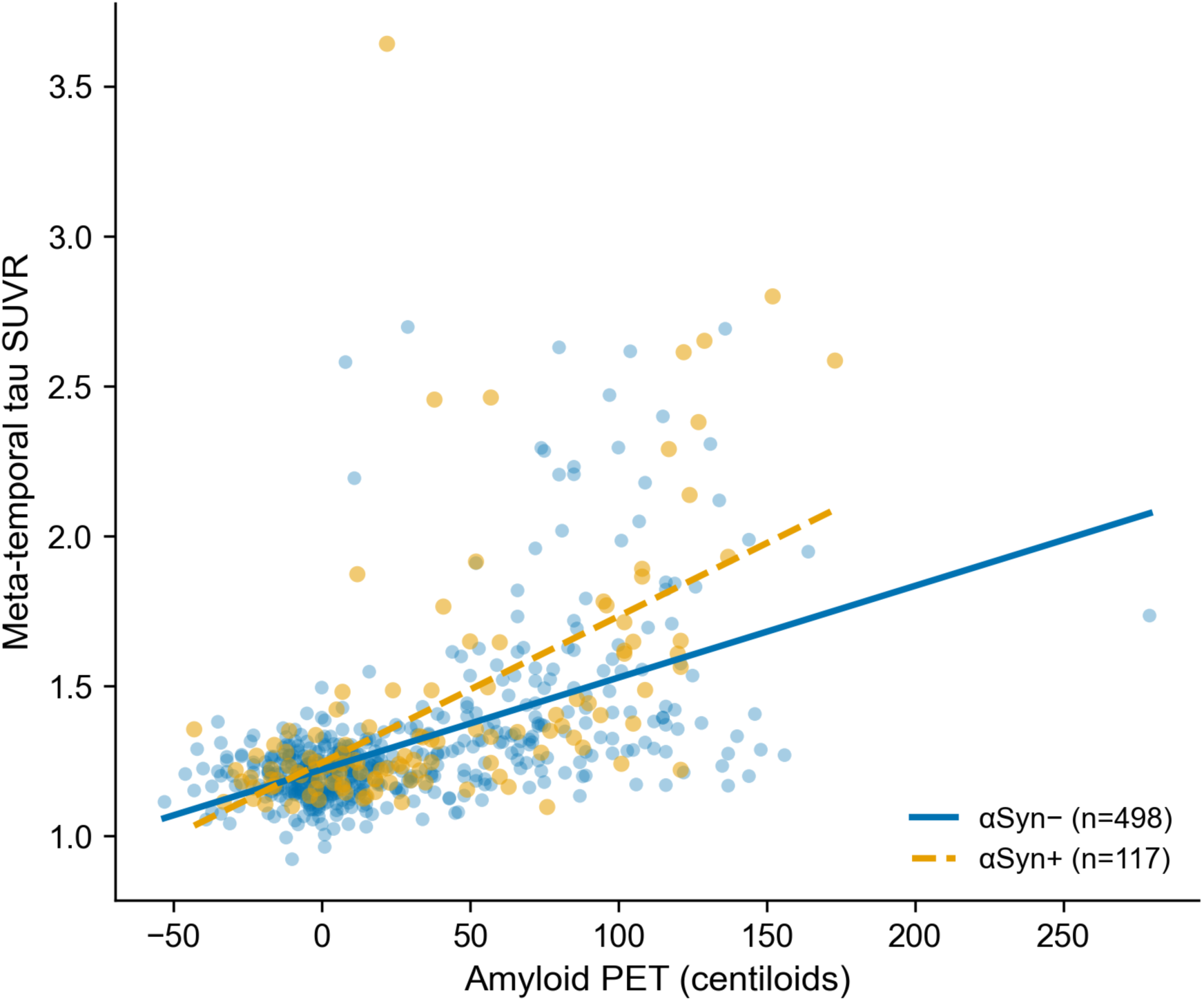
Amyloid burden (centiloids, scan nearest tau PET) and meta-temporal tau PET SUVR by αSyn SAA status (n = 615; 498 αSyn-negative, solid line; 117 αSyn-positive, dashed line). Lines are unadjusted linear fits. The steeper slope in αSyn-positive participants corresponds to the αSyn × centiloids interaction in the adjusted model (β = 0.258, robust 95% CI 0.023–0.493, p = 0.032), which did not survive false discovery rate correction. Abbreviations: αSyn, alpha-synuclein; SAA, seed amplification assay; SUVR, standardized uptake value ratio.

### No detected αSyn modification of the cross-sectional tau–cognition association

In contrast, no αSyn modification of the association between tau and cognition was detected in any domain. Higher meta-temporal tau was associated with lower scores on every cognitive composite (memory: β = −0.440, p < 0.001; executive function: β = −0.434, p < 0.001; language: β = −0.425, p < 0.001; visuospatial: β = −0.366, p < 0.001). The αSyn × tau interaction was not significant for memory (β = −0.038, 95% CI −0.194 to 0.118, p = 0.633), executive function (β = 0.020, 95% CI −0.146 to 0.186, p = 0.811), language (β = 0.003, 95% CI −0.156 to 0.162, p = 0.972), or visuospatial function (β = 0.116, 95% CI −0.094 to 0.325, p = 0.278) (Table 4, Figure 4). At average tau burden, αSyn positivity was associated with lower memory (β = −0.180, p = 0.033), executive function (β = −0.247, p = 0.008), and language (β = −0.187, p = 0.038) scores, with no evidence that these differences depended on tau level. After FDR correction across the 4 domains, the executive function association remained (q = 0.032), whereas the memory and language associations narrowly did not (q = 0.0503 for both); all 3 were attenuated after additional adjustment for diagnosis, CDR-SB, and MMSE, which are themselves cognitive and functional measures, although the executive function difference persisted (β = −0.164, p = 0.039; Supplementary Table S4). These are differences in cognitive level; αSyn positivity was not associated with faster cognitive decline (Supplementary Table S8). With Braak stage-specific tau in place of meta-temporal tau, 1 of 20 αSyn × tau interactions reached nominal significance (memory with Braak I–II tau, β = −0.163, 95% CI −0.316 to −0.009, p = 0.038), which did not survive FDR correction across the 20 tests (q = 0.757; Supplementary Table S2).

**Table 4.** Cross-sectional αSyn × tau interaction on domain-specific cognition (n = 615). Each row is a linear regression of a z-standardized PHC cognitive composite on z-standardized meta-temporal tau SUVR, αSyn status, and their interaction, adjusted for centiloids, age, sex, education, and APOE ε4 carrier status, with heteroscedasticity-consistent (HC3) standard errors.

| Cognitive domain | $\beta$ tau | $\beta$ $\alpha\text{Syn}$ | $\beta$ interaction (95% CI) | p interaction | Adj. $R^2$ |
| --- | --- | --- | --- | --- | --- |
| Memory | -0.440 ($p < 0.001$) | -0.180 ($p = 0.033$) | -0.038 (-0.194, 0.118) | 0.633 | 0.428 |
| Executive function | -0.434 ($p < 0.001$) | -0.247 ($p = 0.008$) | 0.020 (-0.146, 0.186) | 0.811 | 0.322 |
| Language | -0.425 ($p < 0.001$) | -0.187 ($p = 0.038$) | 0.003 (-0.156, 0.162) | 0.972 | 0.289 |
| Visuospatial | -0.366 ($p < 0.001$) | -0.050 ($p = 0.651$) | 0.116 (-0.094, 0.325) | 0.278 | 0.117 |

$\beta$ values are standardized coefficients from linear regression of each z-scored PHC cognitive composite on z-scored meta-temporal tau SUVR, $\alpha\text{Syn}$ status and their interaction, adjusted for centiloids, age, sex, education and APOE $\epsilon 4$ carrier status ( $n = 615$ ); p values and 95% CIs use heteroscedasticity-consistent (HC3) standard errors. Abbreviations: $\alpha\text{Syn}$ , alpha-synuclein; Adj., adjusted; APOE, apolipoprotein E; CI, confidence interval; PHC, Phenotype Harmonization Consortium; SUVR, standardized uptake value ratio.

**Figure 4.**
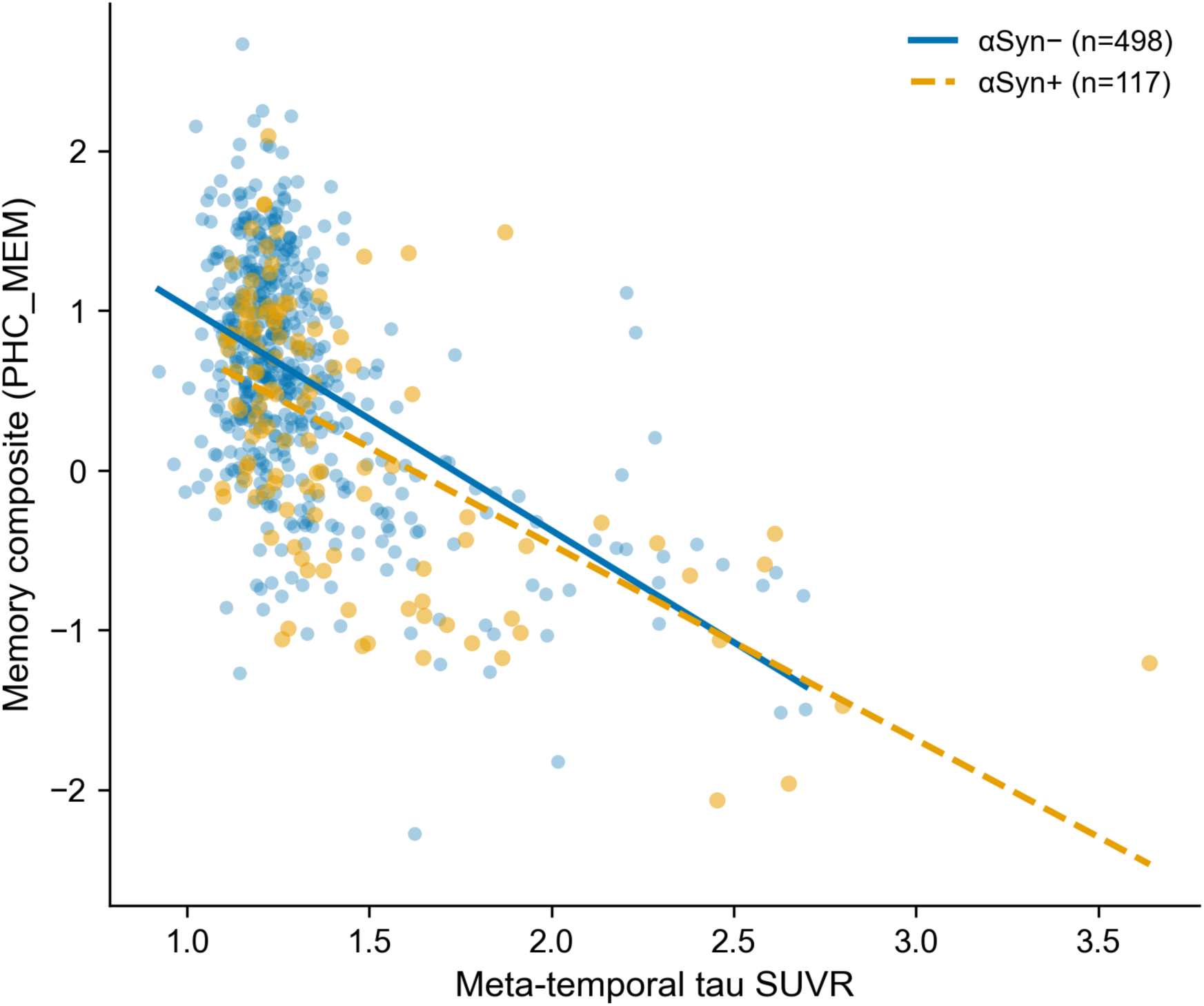
Meta-temporal tau PET SUVR and the PHC memory composite by αSyn SAA status (n = 615). Near-parallel slopes indicate no αSyn × tau interaction on memory in the adjusted model (β = −0.038, robust 95% CI −0.194 to 0.118, p = 0.633); the vertical offset reflects lower memory scores in αSyn-positive participants. Abbreviations: αSyn, alpha-synuclein; PHC, Phenotype Harmonization Consortium; SAA, seed amplification assay; SUVR, standardized uptake value ratio.

### Summary of interaction effects

Figure 5 summarizes the αSyn interaction effects at both steps of the cascade. All 6 amyloid-to-tau interaction coefficients were positive, with robust confidence intervals that excluded zero for meta-temporal and Braak I–IV tau, whereas the 4 tau-to-cognition coefficients were small, with confidence intervals that included zero. In unadjusted regional means (Figure 6), the A+/αSyn+ group had the highest tau SUVR in every Braak region, above the A+/αSyn− group, whereas tau SUVR was low in amyloid-negative participants regardless of αSyn status.

**Figure 5.**
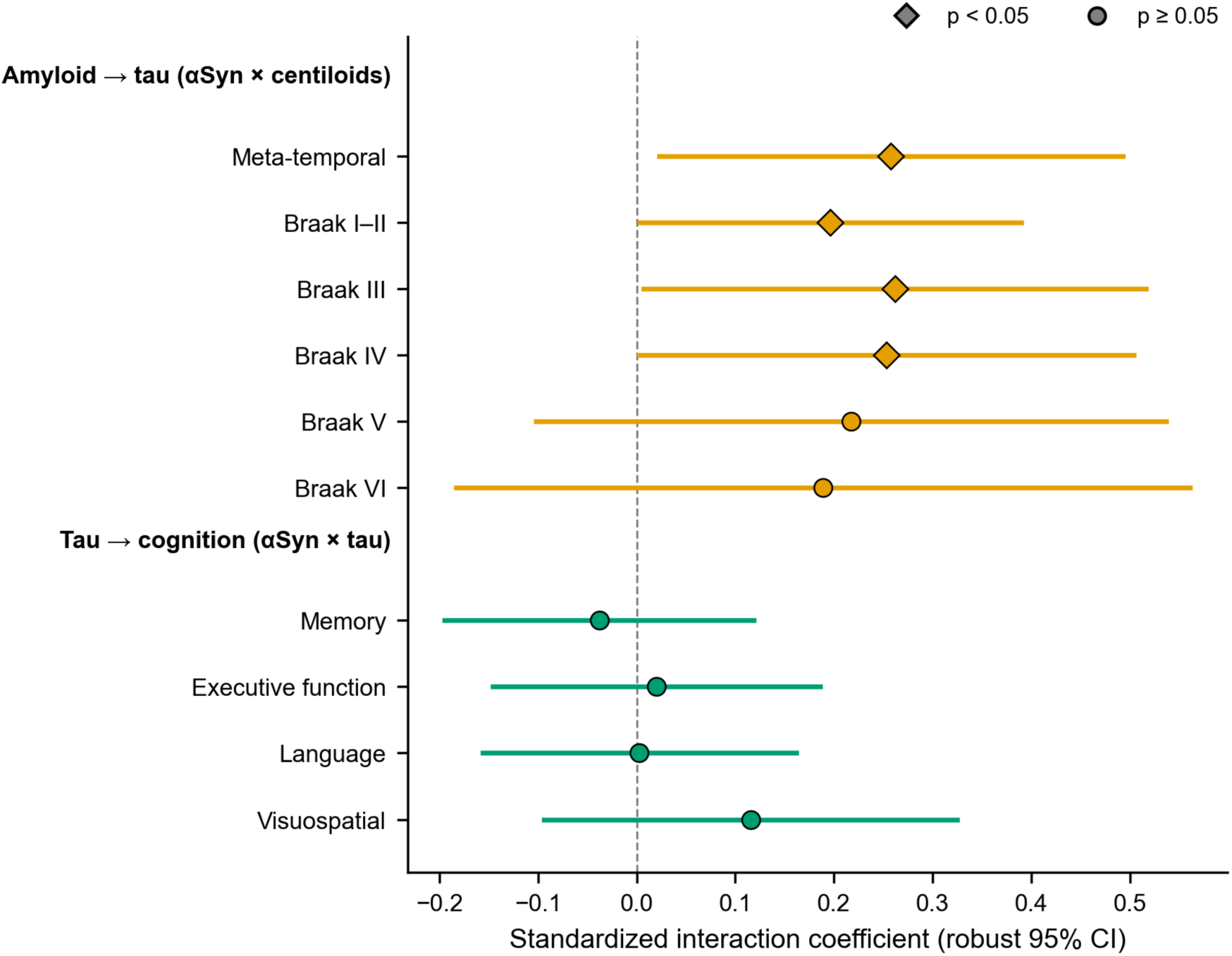
Standardized αSyn interaction coefficients at 2 steps of the AD cascade (n = 615). Upper panel: αSyn × centiloids interaction on regional tau PET SUVR (amyloid-to-tau step). Lower panel: αSyn × meta-temporal tau interaction on each cognitive composite (tau-to-cognition step). Diamonds, p < 0.05; circles, p ≥ 0.05; horizontal bars, heteroscedasticity-consistent 95% CIs. No amyloid-to-tau interaction survived false discovery rate correction. Abbreviations: αSyn, alpha-synuclein; CI, confidence interval; SUVR, standardized uptake value ratio.

**Figure 6.**
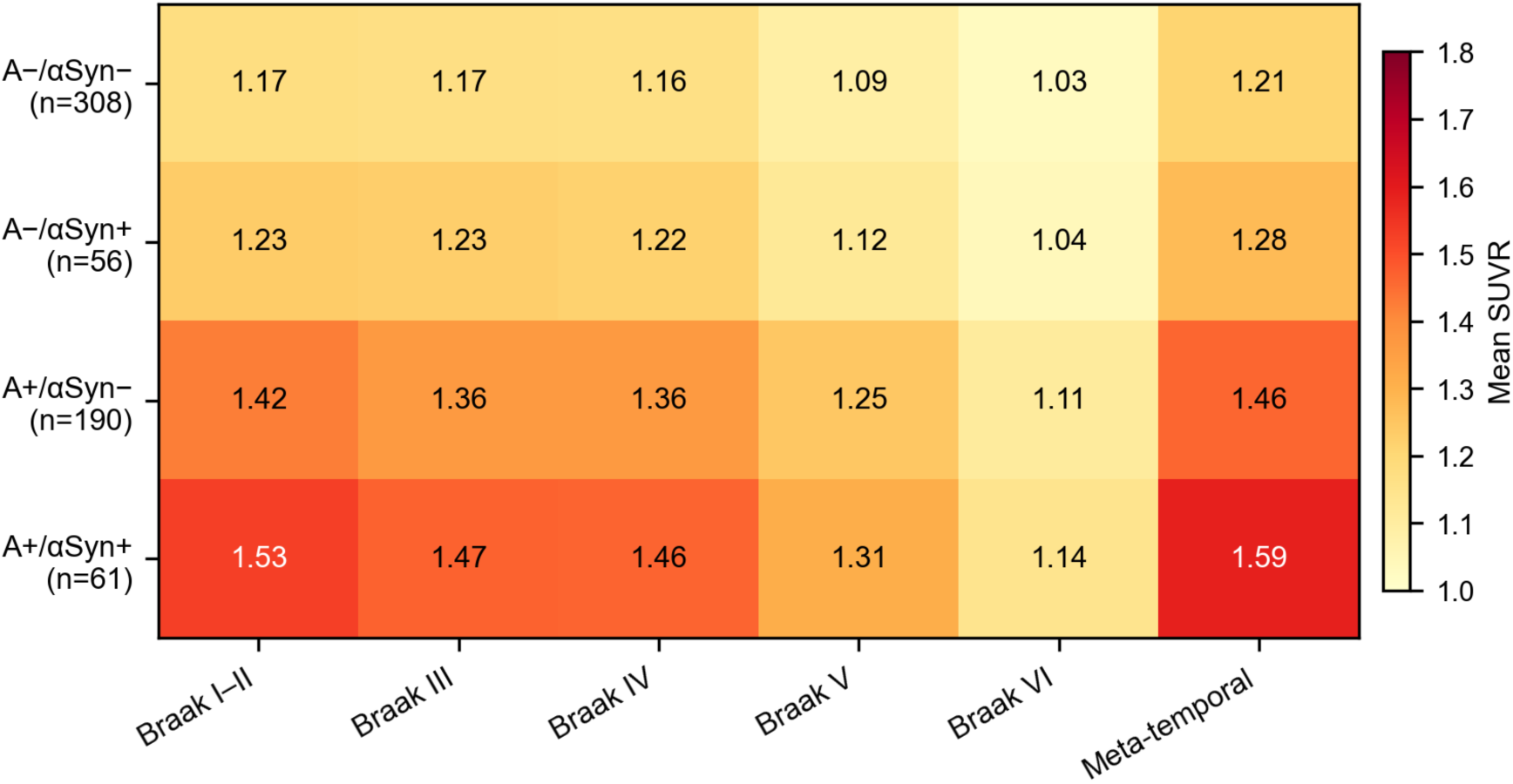
Unadjusted mean regional tau PET SUVR by amyloid and αSyn status (n = 615). Cell values are group means on a yellow-to-red scale; the A+/αSyn+ group had the highest mean SUVR in every Braak region. Abbreviations: A, amyloid (PHC tracer-specific status of the scan nearest tau PET); αSyn, alpha-synuclein; SUVR, standardized uptake value ratio.

### Sensitivity analyses

We conducted 8 sensitivity analyses (4 planned with the primary analysis, 1 added at a coauthor’s request, and 3 added during revision), all with robust standard errors. First, restricting to amyloid-positive participants (n = 251; 61 αSyn-positive), the αSyn × centiloids interaction was larger (meta-temporal β = 0.548, 95% CI 0.093 to 1.003, p = 0.018) and was nominally significant for Braak I–IV but not Braak V–VI. Second, diagnosis-stratified analyses in CN (n = 349) and MCI (n = 200) subgroups showed small, non-significant interactions (CN meta-temporal β = 0.089, p = 0.430; MCI β = 0.112, p = 0.602). Third, in logistic models comparing each AT profile with A−T−, αSyn positivity was not significantly associated with profile membership after adjustment (A−T+ OR = 2.06, 95% CI 0.86 to 4.93, p = 0.104; A+T+ OR = 1.67, 95% CI 0.97 to 2.86, p = 0.062; A+T− OR = 1.13, 95% CI 0.59 to 2.15, p = 0.707). Fourth, in models without the interaction term, αSyn positivity was associated with higher tau in every region except Braak VI (meta-temporal β = 0.318, p = 0.006; Braak III β = 0.340, p = 0.003; Braak VI p = 0.055). Fifth, additional adjustment for clinical diagnosis, CDR-SB, and MMSE reduced the meta-temporal interaction from 0.258 to 0.114 (95% CI −0.117 to 0.346, p = 0.333), and no regional interaction remained significant (all p > 0.33; Supplementary Table S3); the αSyn × tau interactions on cognition also remained non-significant (all p > 0.16; Supplementary Table S4). Sixth, using the earliest examination of every modality in all 636 participants gave the same pattern (meta-temporal β = 0.259, 95% CI 0.034 to 0.484, p = 0.024; nominally significant for Braak I–IV but not V–VI; all q ≥ 0.070), with no αSyn × tau interaction on cognition (all p > 0.30; Supplementary Table S6). Seventh, requiring the CSF sample within 2 years of tau PET (n = 356) gave a larger meta-temporal estimate with wider uncertainty (β = 0.338, 95% CI −0.003 to 0.679, p = 0.052; Supplementary Table S6). Eighth, excluding the 2 Detected-2 participants left the meta-temporal estimate unchanged (β = 0.255, 95% CI 0.020 to 0.491, p = 0.034; Supplementary Table S10). With age taken at the first amyloid PET visit, as in earlier versions of this analysis, the meta-temporal estimate was 0.262 (95% CI 0.027 to 0.497, p = 0.029; Supplementary Table S10).

### Longitudinal tau PET

Of the 636 participants, 341 (68 αSyn-positive) had at least 2 tau PET scans (904 scans; median 2, range 2 to 6), with a median follow-up of 3.5 years (IQR 2.0 to 4.1). Higher baseline amyloid was associated with faster tau accumulation in every region (meta-temporal time × centiloids β = 0.066 SD per year, 95% CI 0.051 to 0.082, p < 0.001). No αSyn modification of this association was detected in any region (meta-temporal time × centiloids × αSyn β = −0.012, 95% CI −0.046 to 0.023, p = 0.513; all regional p > 0.12, all q > 0.59), and αSyn-positive participants did not accumulate tau faster at average amyloid burden (β = 0.016, 95% CI −0.015 to 0.047, p = 0.321) (Supplementary Table S7, Supplementary Figure S1A). At 100 centiloids, model-implied meta-temporal accumulation was 0.050 SUVR per year (95% CI 0.039 to 0.061) in αSyn-negative and 0.050 SUVR per year (95% CI 0.031 to 0.068) in αSyn-positive participants, and the αSyn-positive minus αSyn-negative difference was close to zero at 0, 50, and 100 centiloids (0.007, 0.003, and −0.0004 SUVR per year; all p > 0.23; Supplementary Table S13).

The result held in amyloid-positive participants (n = 146, 38 αSyn-positive; meta-temporal time × αSyn β = 0.009, 95% CI −0.059 to 0.077, p = 0.788; ≥20 centiloids: β = −0.001, p = 0.975), after adjustment for baseline diagnosis (time × centiloids × αSyn β = −0.017, 95% CI −0.051 to 0.018, p = 0.343), and in inverse probability-weighted slope models (β = −0.039, 95% CI −0.140 to 0.062, p = 0.449) (Supplementary Table S10). Participants with tau PET follow-up had higher memory scores at baseline tau PET (0.59 vs 0.40, p = 0.001), were more often APOE ε4 carriers (43.7% vs 33.9%, p = 0.015), and were more often cognitively normal (p = 0.013) than those without, whereas age (73.3 vs 74.4 years, p = 0.082) and αSyn positivity were similar (19.9% vs 18.0%, p = 0.595) (Supplementary Table S9).

Restricting the analysis to participants whose CSF sample was collected no later than 1 year after baseline tau PET (n = 191; 36 αSyn-positive) or within 2 years of it (n = 169; 26 αSyn-positive) did not reveal an interaction (meta-temporal β = −0.033, 95% CI −0.091 to 0.024, p = 0.255; β = −0.046, 95% CI −0.108 to 0.016, p = 0.149; Supplementary Table S13). With the 2-stage approach of Franzmeier et al., the standardized αSyn × centiloids estimate for meta-temporal accumulation was positive (β = 0.261, 95% CI 0.031 to 0.491, p = 0.027 with their covariates; β = 0.195, p = 0.076 with ours) but not significant with robust standard errors (p = 0.123 and p = 0.218; Supplementary Table S12). Participant-specific rates from the first stage correlated with baseline meta-temporal tau (r = 0.96; random intercept–slope correlation 0.92), and the estimate reversed sign when baseline tau was added to the second stage (β = −0.068, p = 0.079) or when unshrunk participant slopes were used (β = −0.254, p = 0.046; robust p = 0.439).

In the exploratory sex analysis, the time × αSyn × sex interaction was significant for meta-temporal tau (β = 0.102 SD per year, 95% CI 0.041 to 0.164, p = 0.001). The 29 αSyn-positive women accumulated 0.042 SUVR per year (95% CI 0.028 to 0.056) and αSyn-positive men 0.011 (95% CI −0.00004 to 0.023), compared with 0.018 in both αSyn-negative women and men. The estimate was similar without adjustment for amyloid (β = 0.100, p = 0.004) and absent for Braak I–II tau (p = 0.884) (Supplementary Table S14).

### Longitudinal cognition

Of the 636 participants, 516 (100 αSyn-positive) had at least 2 cognitive assessments from 6 months before baseline tau PET onward (1,583 assessments; median follow-up 3.2 years, IQR 2.0 to 4.0). Higher baseline meta-temporal tau predicted faster decline in every domain (time × tau β from −0.092 to −0.149 SD per year, all p < 0.001). The time × tau × αSyn interaction was not significant for memory (β = 0.016, 95% CI −0.032 to 0.064, p = 0.520), language (β = 0.020, 95% CI −0.049 to 0.090, p = 0.569), or visuospatial function (β = 0.051, 95% CI −0.029 to 0.131, p = 0.210). For executive function, the interaction was positive (β = 0.082, 95% CI 0.019 to 0.145, p = 0.010, q = 0.042), indicating less steep, not steeper, tau-related executive decline in αSyn-positive participants. This interaction survived FDR correction across the 4 domains, but it was not anticipated, runs opposite to synergy, and has no cross-sectional counterpart, so it requires replication. αSyn positivity was not associated with faster decline at average tau burden in any domain (all p > 0.17) (Supplementary Table S8, Supplementary Figure S1B).

## 4. Discussion

In ADNI participants with amyloid PET, cognition, age, and diagnosis matched to tau PET, αSyn SAA positivity was associated with a modestly steeper cross-sectional association between amyloid and tau PET in the meta-temporal region and Braak I–IV, but this interaction did not survive correction for multiple comparisons and was attenuated to non-significance after adjustment for clinical diagnosis and severity. Over a median of 3.5 years of repeated tau PET, we detected no αSyn-related modification of amyloid-related tau accumulation, although an exploratory analysis suggested faster accumulation in αSyn-positive women. We detected no αSyn modification of the cross-sectional association between tau and cognition; over time, tau-related executive decline was less steep in αSyn-positive participants, a finding opposite in direction to synergy that requires replication. The data do not support αSyn-driven acceleration of amyloid-related tau spread over a 3- to 4-year window, although they cannot exclude a moderate effect.

These results partly replicate and partly diverge from those of Franzmeier et al., who analyzed an overlapping ADNI sample [16]. The direction of the cross-sectional findings agrees: both studies found a steeper amyloid–tau association in αSyn-positive participants, and our largest estimates in Braak III–IV, the stages at which tau extends from medial temporal structures into the neocortex [29–31], fit neuropathological evidence of greater neocortical tau burden in AD patients with Lewy body co-pathology [6]. The strength of the cross-sectional evidence, however, depended on the analysis: with heteroscedasticity-consistent standard errors, the interaction was nominally significant in 4 of 6 regions, did not survive FDR correction, and lost more than half its size after adjustment for clinical stage. The longitudinal findings do not agree. Franzmeier et al. reported faster amyloid-related tau accumulation in 290 participants followed for about 2.5 years; with 341 participants, a median of 3.5 years, and 1-stage mixed-effects models, we detected no αSyn × amyloid × time interaction, although the confidence interval (upper limit 0.023 SD per year per SD of centiloids) does not exclude a small positive effect. In our data, the analytic approach reproduced the discrepancy. Their 2-stage approach, applied to our participants and scans, gave a positive estimate (β = 0.261 with their covariates, versus 0.175 in their study), but that estimate depended on baseline tau. Change rates predicted from a mixed model without covariates borrow strength from each participant’s baseline value, 186 of our 341 participants had only 2 scans, and the estimate reversed sign once baseline tau was accounted for. The 1-stage model avoids this dependence by estimating change and its predictors jointly [27]. Differences in image processing (Schaefer-atlas SUVRs in their study, PHC composites in ours), in the window allowed between modalities, and in follow-up length could also contribute (Supplementary Table S11), and neither study has enough αSyn-positive participants with repeated scans (68 here) to settle the question.

The exploratory sex analysis agrees with Mak et al., who found faster tau accumulation in SAA-positive women than in SAA-positive men in a partly overlapping ADNI sample [28]. In our data the association persisted after adjustment for amyloid, which that study did not include as a covariate, and was present for meta-temporal tau, the composite closest to the meta-temporal region they analyzed, but not for the narrower Braak I–II region. It was not prespecified, rests on 29 αSyn-positive women, and comes from overlapping data, so it needs testing in an independent cohort before it informs sex-specific trial enrichment. It also shows that a null αSyn × amyloid × time term does not rule out αSyn-related differences in tau accumulation within subgroups.

Several preclinical mechanisms could link αSyn to tau. Distinct αSyn fibril strains cross-seed tau aggregation in primary neurons and in vivo [8], Aβ-seeded tau fibrils act as potent templates for prion-like tau propagation [32], and soluble αSyn oligomers alter the aggregation kinetics of Aβ and tau [9, 10], while all three proteins share prion-like propagation pathways [33]. Triple-transgenic mice expressing Aβ, tau, and αSyn develop more severe pathology and cognitive impairment than mice with one or two of these proteinopathies [7]. These mechanisms predict faster tau accumulation in αSyn-positive individuals, which our longitudinal data did not show. A weak cross-sectional interaction that depends on clinical stage and has no detected longitudinal counterpart does not support αSyn as a driver of amyloid-related tau spread; whether αSyn seeding arises later in the amyloid-to-tau sequence cannot be determined from these observational data.

Cross-sectionally, the αSyn × tau interaction on cognition was not significant in any domain (all interaction p > 0.27), reached nominal significance for only 1 of 20 Braak-stage tau measures, and was not significant after diagnosis adjustment. Longitudinally, no αSyn modification of tau-related decline was detected for memory, language, or visuospatial function, although the confidence intervals do not establish equivalence. The executive function interaction survived correction within its family but ran opposite to synergy and had no cross-sectional counterpart; it may reflect a true difference, for example a floor effect in participants already impaired at baseline, or chance, and it requires replication. Franzmeier et al. likewise found no αSyn × tau PET interaction on global cognitive decline [16]. The main effects point elsewhere: at mean tau, αSyn positivity was associated with lower executive function scores, with weaker associations for memory and language, which is compatible with contributions that tau PET does not capture, such as synaptic dysfunction, dopaminergic deficits, or cortical Lewy body pathology [34], although cross-sectional associations cannot show that these contributions are independent of tau. Associations of αSyn with cognition or metabolism beyond measured AD pathology have been reported in clinically unimpaired adults [15], in amnestic patients with AD co-pathology [17], in posterior hypometabolism after accounting for CSF amyloid and tau [35], and in dementia with Lewy bodies with AD co-pathology [36]. For clinicians, these data do not show that tau PET carries different prognostic information in αSyn-positive and αSyn-negative patients, but they do not establish that it carries the same information either; a positive SAA was associated with lower executive performance at the average tau level, and we did not detect faster decline over a median of 3.2 years.

These findings bear on anti-amyloid therapy. Lecanemab slowed, but did not halt, clinical decline in early AD [37]. αSyn co-pathology is one plausible reason why treated patients continue to decline, and αSyn-positive patients have been proposed to benefit most from early amyloid removal if αSyn accelerates the amyloid-to-tau transition [16]. Our longitudinal data did not detect overall acceleration, so the more defensible use of αSyn SAA status in trials is as a stratification variable for cognitive outcomes, because αSyn positivity was associated with poorer executive function at the average tau level here and with faster cognitive decline in a larger ADNI analysis [13], rather than as a marker of expected benefit. Whether αSyn status modifies treatment response can be answered only in randomized cohorts, for example by prespecifying αSyn SAA status as a stratifier in anti-amyloid trials and testing a treatment × αSyn interaction on the primary clinical endpoint [38]. Tau-clinical mismatch, in which patients decline faster than their tau burden predicts and more often carry αSyn co-pathology, offers a complementary route to the same stratification [39].

Three alternative explanations deserve consideration. First, αSyn co-pathology may be more common in individuals already further along the amyloid-to-tau sequence, rather than causing tau accumulation; a cross-sectional interaction that weakens after adjustment for clinical stage and has no detected longitudinal counterpart fits this explanation, as does the rise in SAA positivity with age and disease stage in ADNI [12]. Second, αSyn SAA positivity may represent a marker of broader cellular dysfunction, including neuroinflammation, lysosomal impairment, or generalized proteostatic failure, rather than a specific αSyn-mediated effect. Third, residual confounding by unmeasured co-pathologies cannot be excluded; TDP-43 pathology [40] and cerebrovascular disease are common in older adults and could contribute to both SAA positivity and tau burden. Serial SAA and tau PET within ADNI [18] would allow these explanations to be separated.

Three features argue against over-interpreting the cross-sectional interaction, and one argues for it. First, its statistical support depended on the variance model: conventional standard errors gave p = 0.002 to 0.034 in all 6 regions, whereas heteroscedasticity-consistent standard errors, which the residual structure required, gave p = 0.032 to 0.048 in 4 regions and no interaction that survived FDR correction. Second, adjustment for diagnosis, CDR-SB, and MMSE reduced the meta-temporal estimate from 0.258 to 0.114; because these variables lie partly downstream of tau pathology, this adjustment may remove true signal [41] and answers a different conditional question rather than correcting the primary estimate, but it also shows that the interaction is entangled with clinical stage. Third, the diagnosis-stratified estimates were small. In its favor, the estimate itself was stable across samples (0.255 to 0.338 in the earliest-examination, CSF-restricted, and Detected-2-excluded analyses) and larger in amyloid-positive participants, so the uncertainty lies in its precision rather than its direction. The lack of detectable αSyn-related modification of the tau–cognition relationship was consistent across Braak-stage tau measures, diagnosis-adjusted models, and alternative samples, and held longitudinally in 3 of 4 domains.

## Strengths and limitations

This study combines CSF αSyn SAA, amyloid PET, regional tau PET spanning the full Braak hierarchy, structural MRI, APOE genotyping, domain-specific cognitive composites, and repeated tau PET over a median of 3.5 years. The harmonized PHC pipeline provides a processing stream independent of that used by Franzmeier et al., which makes the comparison with their results informative. Amyloid PET and cognition were matched to within 1 year of tau PET, age and diagnosis were taken at tau PET, inference used standard errors robust to non-constant variance, the longitudinal estimate was compared with the earlier ADNI approach in the same participants, and the cross-sectional tau–cognition findings were consistent across sensitivity analyses.

Several limitations apply. First, amyloid PET, cognition, age, and diagnosis were matched to tau PET, but MRI was more than 1 year from tau PET in 41% of the primary sample, and CSF was collected a median of 1.73 years from tau PET, after the tau PET date in more than half of participants. SAA status was measured once and treated as time-invariant, so status at baseline and during follow-up may be misclassified, with an uncertain effect on interaction estimates, and a result obtained after baseline is not a prospectively available marker; restricting the longitudinal analysis by CSF collection date did not change the result. Second, requiring every modality selected participants able to complete a multimodal protocol, and participants with tau PET follow-up were less impaired than those without, although weighting for the probability of follow-up did not change the result. Third, only 68 αSyn-positive participants had repeated tau PET, and 38 of them were amyloid-positive, so moderate longitudinal effects cannot be excluded, and the exploratory sex analysis rests on 29 αSyn-positive women. Fourth, the SAA is binary and detects neocortical Lewy pathology more reliably than limbic or amygdala-predominant disease [12], so early αSyn co-pathology was likely underdetected. Fifth, flortaucipir off-target binding and spill-in from the choroid plexus and white matter affect Braak I–II and hippocampal estimates [42]. Sixth, the diagnosis-stratified analyses were underpowered, and the dementia subgroup was small (n = 66). Seventh, neurodegeneration status was defined by a cohort-median split of SPARE-AD rather than an external cut point, so the ATN profile frequencies are specific to this sample. Finally, ADNI participants are predominantly non-Hispanic White and highly educated, which may limit the generalizability of these findings to more diverse populations.

## Conclusion

αSyn co-pathology was associated with a modestly steeper cross-sectional coupling between amyloid and tau, but the interaction was sensitive to correction for multiple testing and to adjustment for clinical stage, no αSyn-related acceleration of amyloid-related tau accumulation was detected over a median of 3.5 years, and no αSyn modification of the cross-sectional association between tau and cognition was detected; longitudinally, tau-related executive decline was unexpectedly less steep in αSyn-positive participants and needs replication.

αSyn positivity may mark patients further along the AD continuum, and it was associated with poorer executive function after adjustment for measured tau. Faster tau accumulation in αSyn-positive women, seen here in an exploratory analysis, needs confirmation in an independent cohort. Longer tau PET follow-up in αSyn-positive participants, and prespecified αSyn stratification in anti-amyloid trials, are needed before αSyn status guides treatment decisions.

## Statements and declarations

### Ethics approval and consent to participate

Data used in this study were obtained from the Alzheimer’s Disease Neuroimaging Initiative (ADNI) database. The ADNI study was approved by the Institutional Review Board or Independent Ethics Committee at each participating site, and all participants provided written informed consent in accordance with the Declaration of Helsinki. This study represents a secondary analysis of de-identified ADNI data conducted under the ADNI Data Use Agreement.

### Availability of data and materials

The data used in this study were obtained from the Alzheimer’s Disease Neuroimaging Initiative (ADNI) database. ADNI data are publicly available to qualified investigators upon registration and approval at https://adni.loni.usc.edu.

### CRediT authorship contribution statement

A.N.: Conceptualization, methodology, validation, investigation, resources, supervision, project administration, writing (original draft preparation), writing (review and editing), J.P.R.: Methodology, formal analysis, data curation, visualization, investigation, writing (review and editing). Funding acquisition: Not applicable. All authors have read and approved the final manuscript.

### AI Disclosure

Portions of the statistical code were developed with the assistance of an AI model for code development and debugging (Anthropic Claude Code and Claude Science Opus 4.8, 5.0, and 5.5), and all analyses were independently replicated and reproduced by the authors in R (version 4.5.2) and Stata/MP (version 18; macOS, Apple silicon). Full analysis scripts in these 3 platforms are available upon request. The authors are responsible for study conception, design, variable selection, data interpretation, and verification of all analytical outputs and scientific claims.

## Acknowledgments

Data collection and sharing for this project was funded by the Alzheimer’s Disease Neuroimaging Initiative (ADNI) (National Institutes of Health Grant U01 AG024904) and DOD ADNI (Department of Defense award number W81XWH-12-2-0012). ADNI is funded by the National Institute on Aging, the National Institute of Biomedical Imaging and Bioengineering, and through generous contributions from the following: AbbVie, Alzheimer’s Association; Alzheimer’s Drug Discovery Foundation; Araclon Biotech; BioClinica, Inc.; Biogen; Bristol-Myers Squibb Company; CereSpir, Inc.; Cogstate; Eisai Inc.; Elan Pharmaceuticals, Inc.; Eli Lilly and Company; EuroImmun; F. Hoffmann-La Roche Ltd and its affiliated company Genentech, Inc.; Fujirebio; GE Healthcare; IXICO Ltd.; Janssen Alzheimer Immunotherapy Research & Development, LLC.; Johnson & Johnson Pharmaceutical Research & Development LLC.; Lumosity; Lundbeck; Merck & Co., Inc.; Meso Scale Diagnostics, LLC.; NeuroRx Research; Neurotrack Technologies; Novartis Pharmaceuticals Corporation; Pfizer Inc.; Piramal Imaging; Servier; Takeda Pharmaceutical Company; and Transition Therapeutics. The Canadian Institutes of Health Research is providing funds to support ADNI clinical sites in Canada. Private sector contributions are facilitated by the Foundation for the National Institutes of Health (www.fnih.org). The grantee organization is the Northern California Institute for Research and Education, and the study is coordinated by the Alzheimer’s Therapeutic Research Institute at the University of Southern California. ADNI data are disseminated by the Laboratory for Neuro Imaging at the University of Southern California.

## Abbreviations

The following abbreviations are used in this manuscript:

AD: Alzheimer’s disease
APOE: Apolipoprotein E
ATN: Amyloid-Tau-Neurodegeneration
Aβ: Amyloid-beta
CDR-SB: Clinical Dementia Rating Sum of Boxes
CI: Confidence interval
CN: Cognitively normal
CSF: Cerebrospinal fluid
DLB: Dementia with Lewy bodies
FDR: False discovery rate
IQR: Interquartile range
MCI: Mild cognitive impairment
MMSE: Mini-Mental State Examination
MRI: Magnetic resonance imaging
NIA-AA: National Institute on Aging–Alzheimer’s Association
OLS: Ordinary least squares
PET: Positron emission tomography
PHC: Phenotype Harmonization Consortium
PPMI: Parkinson’s Progression Markers Initiative
SAA: Seed amplification assay
SD: Standard deviation
SPARE-AD: Spatial Pattern of Abnormality for Recognition of Early Alzheimer’s Disease
SUVR: Standardized uptake value ratio
αSyn: α-Synuclein

## Supplementary material

### Supplementary table legends

Table S1. False discovery rate correction of the amyloid–tau interaction analyses.

Table S2. Braak-stage-specific tau × αSyn interaction analyses for cognitive outcomes.

Table S3. Amyloid-tau interaction models additionally adjusted for diagnosis, CDR-SB, and MMSE.

Table S4. Tau–cognition models additionally adjusted for diagnosis, CDR-SB, and MMSE: αSyn × tau interaction and αSyn main effect.

Table S5. Intervals between baseline tau PET and the other modalities in the primary sample (n = 615) and the earliest-examination cohort (N = 636).

Table S6. Amyloid–tau and tau–cognition interactions in alternative samples: earliest examinations of every modality (N = 636) and CSF sample within 2 years of tau PET (n = 356).

Table S7. Linear mixed-effects models of longitudinal regional tau PET (n = 341).

Table S8. Linear mixed-effects models of longitudinal cognitive composites (n = 516).

Table S9. Participants observed per year of follow-up and baseline comparison of participants with and without tau PET follow-up.

Table S10. Additional sensitivity analyses: exclusion of Detected-2 results and the earlier age anchor (primary sample, robust standard errors), amyloid-positive and diagnosis-adjusted longitudinal tau models, and inverse probability-weighted slope models.

Table S11. Design of the present study compared with Franzmeier et al. (2025).

Table S12. αSyn × centiloids interaction on tau accumulation from the 1-stage mixed model and from the 2-stage approach of Franzmeier et al. in the same participants.

Table S13. Longitudinal tau models restricted by CSF collection date, and model-based αSyn group differences in annual meta-temporal tau change at 0, 50, and 100 centiloids.

Table S14. Exploratory αSyn × sex × time models of tau accumulation.

### Supplementary figure legend

Supplementary Figure S1. Longitudinal analyses. (A) Model-implied annual change in meta-temporal tau SUVR across baseline amyloid burden in αSyn SAA-negative (blue) and αSyn SAA-positive (red) participants (n = 341), from a linear mixed-effects model with covariates at reference values; shaded bands show 95% CIs, and tick marks show individual baseline centiloid values. Curves are truncated at the 95th percentile of centiloids among αSyn-positive participants. (B) Time × baseline meta-temporal tau × αSyn interaction estimates for each cognitive composite (n = 516); bars show 95% CIs, and green indicates p < 0.05. Abbreviations: αSyn, alpha-synuclein; SAA, seed amplification assay; SUVR, standardized uptake value ratio; SD, standard deviation.

